# Transcriptional Profiles of Non-neuronal and Immune Cells in Mouse Trigeminal Ganglia

**DOI:** 10.1101/2023.08.18.553897

**Authors:** Jennifer Mecklenburg, Sergey A. Shein, Anahit H. Hovhannisyan, Yi Zou, Zhao Lai, Shivani Ruparel, Alexei V. Tumanov, Armen N. Akopian

**Affiliations:** Department of Endodontics, School of Dentistry, The University of Texas Health Science Center at San Antonio (UTHSCSA), San Antonio, Texas 78229; Molecular Medicine and Molecular Genetics Departments, School of Medicine, UTHSCSA, San Antonio, Texas 78229; Microbiology, Immunology & Molecular Genetics Departments, School of Medicine, UTHSCSA, San Antonio, Texas 78229; Greehey Children’s Cancer Research Institute, UTHSCSA, San Antonio, Texas 78229

**Author notes:** Corresponding author: Armen N. Akopian, The School of Dentistry, University of Texas Health Science Center @ San Antonio (UTHSCSA) 7703 Floyd Curl Drive, San Antonio, TX 78229-3900.

**Keywords:** Trigeminal ganglia, Glia, Immune cells, Stromal cells, Macrophages, Neutrophils, single-cell RNA-seq

## Abstract

Non-neuronal cells constitute 90-95% of sensory ganglia. These cells play critical roles in modulation of nociceptive signal transmissions by sensory neurons. Accordingly, the aim of this review-study was to identify, profile and summarize TG non-neuronal cell types in naïve male mice using published and our own data generated by single-cell RNA sequencing (scRNA-seq), flow cytometry (FC) and immunohistochemistry (IHC). TG contains 5 types of non-neuronal cells: glial, fibroblasts, smooth muscle, endothelial and immune cells. There is agreement among publications for glial, fibroblasts, smooth muscle and endothelial cells. Based on gene profiles, glial cells were classified as Schwann cells and satellite glial cells (SGC). *Mpz* had dominant expression in Schwann cells, and *Fabp7* is specific for SCG. Two types of Col1a2^+^ fibroblasts located throughout TG were distinguished using gene profiles. TG smooth muscle and endothelial cells representing blood vessels were detected with well recognized markers. Our study split reported single TG immune cell group into 3 types of macrophages and 4 types of neutrophils. Macrophages were located among neuronal bodies and nerve fibers, and were sub-grouped by unique transcriptomic profiles and using *Ccr2*, *Cx3cr1* and *Iba1* as markers. S100a8^+^ neutrophils were located in dura surrounding TG and were sub-grouped by clustering and expressions of *Csf3r*, *Ly6G, Ngp, Elane* and *Mpo*. Overall, generated and summarized here dataset on non-neuronal TG cells could provide essential and fundamental information for studies on cell plasticity, interactomic network between neurons and non-neuronal cells and function during variety of pain conditions in the head and neck region.

## Introduction

Multiple reports assigned critical roles in nociceptive signal transmission to dorsal root ganglion (DRG) glial (Huang et al., 2013;Donnelly et al., 2020;Gazerani, 2021;McGinnis and Ji, 2023) and immune cells, especially macrophages and neutrophils (Ji et al., 2016;Yu et al., 2020;Domoto et al., 2021;Lesnak et al., 2023). It is suggested that ganglion non-neuronal cells are capable of sensitizing neurons (Ji et al., 2016;Donnelly et al., 2020;Gazerani, 2021) by directly communicating with them and changing their gating properties (Domoto et al., 2021;Haberberger et al., 2023;McGinnis and Ji, 2023). Accordingly, information on transcriptional profiles for non-neuronal sensory ganglion cells is critically important. Thus, this fundamental information could be used to explain a variety of mechanisms for interactions of sensory neuron soma with non-neuronal cells within the ganglia. For example, these data could be used for calculating interactomic network between sensory neurons and non-neuronal cells (Wangzhou et al., 2021). This dataset could also constitute baseline in investigation of TG non-neuronal cell plasticity in different pain models and conditions for the head and neck region.

Single-nuclei RNA sequencing (snRNA-seq) of DRG and TG cells have previously generated transcriptomic profiles for both sensory neurons and non-neuronal ganglion cells (Drokhlyansky et al., 2020;Renthal et al., 2020;Sharma et al., 2020;Yang et al., 2022). Single-cell RNA sequencing studies are complex and often produce variable outcomes. These outcomes depend on several factors such as ganglia type (DRG vs TG vs nodose ganglia), single-nuclei versus single-cell RNA-seq, nuclei/cell isolation approaches, sequencing depth and clustering analysis (Drokhlyansky et al., 2020;Renthal et al., 2020;Sharma et al., 2020;Yang et al., 2022). Hence, every independent study adds novel information as well as refine previously reported data. Thus, 7 non-neuronal subtypes, including satellite glia (SGC), myelinating and non-myelinating Schwann cells, Mgp^+^ and Dcn^+^ fibroblasts, immune cells, and vascular endothelial cells, were identified in TG using snRNA-seq (Yang et al., 2022), when nuclei were isolated using density gradient method (Renthal et al., 2020). Other snTNA-seq and single-cell RNA-sequencing (scRNA-seq) studies reported 8 (Chu et al., 2023) or 5 (Sharma et al., 2020) types of non-neuronal cells in TG: SGC, myelinating and non-myelinating Schwann cells, fibroblasts, myofibroblasts, immune cells, and vascular endothelial cells. snRNA-seq of DRG cells revealed 9 types of non-neuronal cells, including SGC, myelinating and non-myelinating Schwann cells, one group of fibroblasts, pericytes, and vascular endothelial cells, as well as three types of immune cells: a group of macrophages, B-cells, and single group of neutrophils (Renthal et al., 2020).

Multiple studies on roles of ganglion glial cells in regulation of sensory neurons requires specific markers distinguishing SGC from Schwann cells as well as from other non-neuronal cells. It is not entirely clear whether such markers exist. Function of ganglion fibroblasts is largely unknown. In this respect, more information on their gene profiles and locations within TG is needed. Reports showed several immune cell types in DRG (Renthal et al., 2020), and only one group in TG (Sharma et al., 2020;Yang et al., 2022;Chu et al., 2023). Again, additional studies regarding this topic could be valuable. Accordingly, the aim of this review-study was to identify, profile and summarize TG non-neuronal cell types in naïve male mice using published and our own data generated by single-cell RNA sequencing (scRNA-seq), flow cytometry (FC) and immunohistochemistry (IHC).

## Materials and Methods

### Ethical Approval and Mouse lines

The reporting in the manuscript follows the recommendations in the ARRIVE guidelines (PLoS Bio 8(6), e1000412,2010). We also followed guidelines issued by the National Institutes of Health (NIH) and the Society for Neuroscience (SfN) to minimize the number of animals used and their suffering. All animal experiments conformed to protocols approved by the University Texas Health Science Center at San Antonio (UTHSCSA) Institutional Animal Care and Use Committee (IACUC). Protocol numbers are 20190114AR and 20220069AR. Experiments were performed on following male mice: 10-18-week-old C57BL/6 wild-type; Col1a2-cre-ER (stock: 029567); tdTomato (aka Ai14; stock: 007914); and the Ccr2^RFP^/Cx3cr1^GFP^ (Stock No: 032127) on the B6.129 background. All mouse lines were purchased from the Jackson Laboratory (Bar Harbor, ME) and were bred in UTHSCSA LAR facilities.

### TG isolation and single-cell preparation

There are several approaches to dissect TG tissues. One of them collect TG with surrounding dura, while other approach leads to isolation of dura-free TG. We have isolated TG with surrounding dura for scRNA-seq, TG without dura for IHC and both preparations of TG with and without dura for FC. Briefly, prior to TG dissections, animals were perfused with cold PBS to eliminate contributions of immune cells from blood to scRNA-seq and FC data. For IHC, mice were perfused with 4% paraformaldehyde prior tissue dissections. TG were dissected from the skull base after removal of the brain. For dura-free TG, V1-3 were cut close to TG, and then dura-free TG was lifted by spatula. For TG with dura, continuous cuts was made all around TG, resulting in dissected TG covered by dura.

Dissected TG were collected in ice-cold HBSS buffer and subjected to preparation of single-cell suspension for scRNA-seq or flow cytometry. Single cell suspension was generated using Liberase and Dispase II as described previously (Mecklenburg et al., 2020). After this step, single cell suspension was processed in two different ways. For the first scRNA-seq experiment, fraction enriched with sensory neurons were obtained using Percoll gradient as described previously (Mecklenburg et al., 2020). For the second scRNA-seq experiment, Helix NP Blue (Bio Legend), which is impermeant to live cells, was used for the discrimination of all live cells. Briefly, single-cell suspension was stained for 10-60 min at room temperature with 100nM Helix NP Blue. Helix NP Blue is a blue-emitting dye with an excitation/emission max of 430 nm/470 nm. Therefore, viable TG cells were FACS sorted prior to scRNA seq using the Brilliant Violet 421™ detection channel, a 70 μm nozzle and a flow rate of 2 on a BD FACSARIA II into a 1.5 mL microcentrifuge tube containing 15 μl of 1X PBS,0.04% BSA, and 0.1 U/ul RNase inhibitor.

For flow cytometry experiments, single-cell suspensions after the Liberase-Dispase step (see above) were stained with a panel of antibodies as described below.

### Single-cell RNA-sequencing (scRNA-seq) procedures, clustering, visualization, and annotation

TG from three mice were used to generate single-cell suspension for one experiment. Experiment was performed in two independent replicates: first preparation used Percoll step (Mecklenburg et al., 2020) and the second used Helix NP Blue (see previous sub-section of the Material and Methods”). All clustering methods, visualization (tSNE, UMAP) and analysis were done by 10X tool loupe browser (https://support.10xgenomics.com/single-cell-gene-expression/software/visualization/latest/what-is-loupe-cell-browser). The 10X single cell raw sequencing data from both Next Generation Sequencing runs were processed following 10X single cell gene expression pipeline (https://www.10xgenomics.com/support/single-cell-gene-expression). 10X software Cell-Ranger mkfastq was used for base-calling and generating raw fastqs. Cell-Ranger count was used to align fastq reads to reference transcriptome (refdata-gex-mm10-2020-A) and to generate gene counts matrix and a “cloupe” file. Then the “loupe file” was put in “Loupe browser” for data visualization and preliminarily analysis. Cells with > 500 unique genes, < 15,000 total UMIs, and < 10% of the counts deriving from mitochondrial genes were included for analysis. We used two similar different clustering approaches. For the first scRNA-seq experiment, K-means was employed (*Fig 1A*), while for the second scRNA-seq experiment, Graph-Based approach was used for analysis (*Fig 1B*). For both runs, we employed tSNE plots for visualization. Commonly, doublet or low-quality clusters could significantly be enriched for at least 4 mitochondrial genes (fold change (FC)>2, false discovery rate (Padj) < 0.05), have no enriched cluster marker genes (FC>5, Padj < 0.05). We did not have such clusters in the final presented here data.

**Figure 1.**
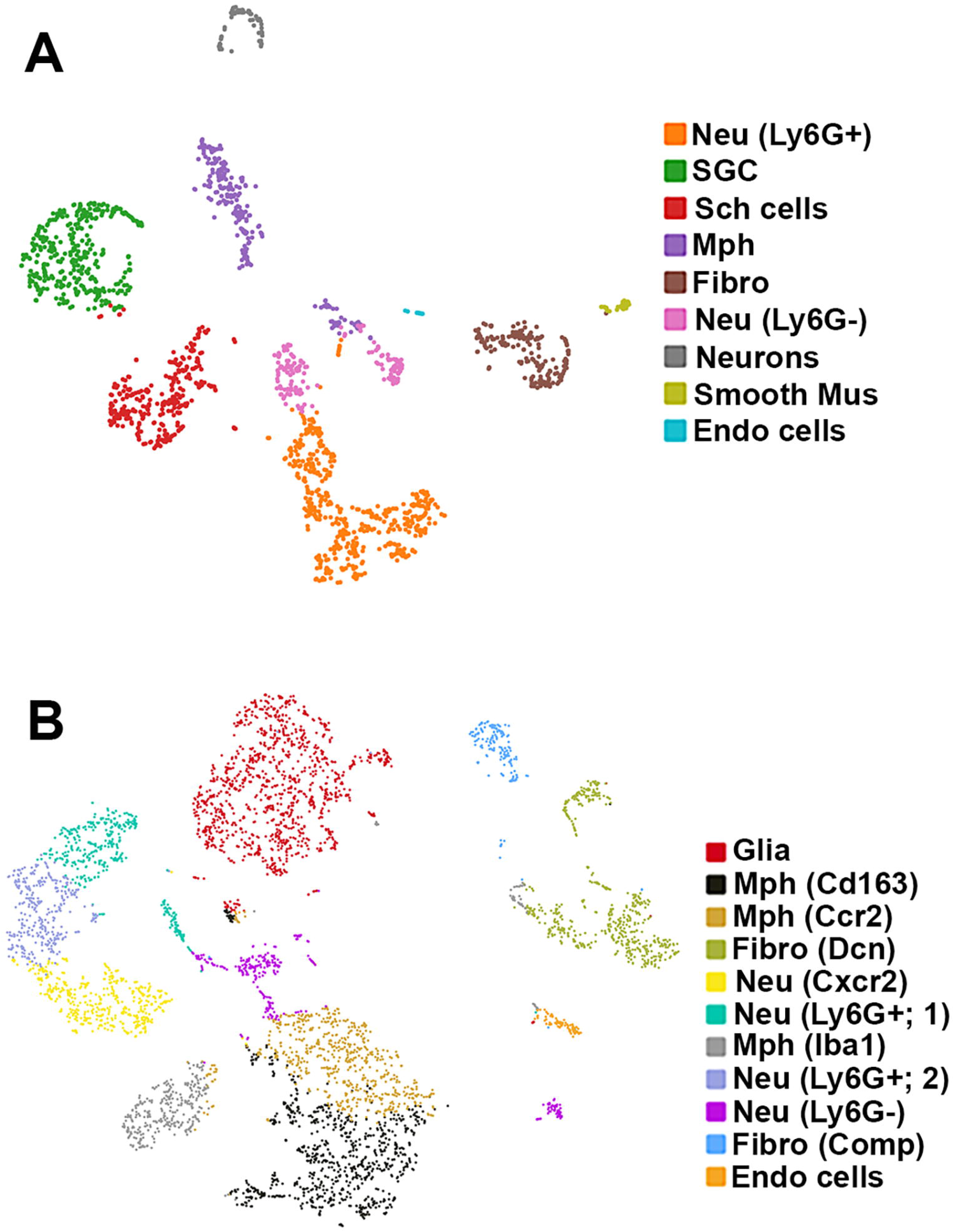
scRNA-seq of mouse TG non-neuronal cells. Two independent replicates were performed. (**A**) tSNE plot of scRNA-seq data from experiment-1 from 2859 live FACS isolated TG cells from 3 mice. (**B**) tSNE plot of scRNA-seq data from experiment-2 from 6362 live FACS isolated TG cells from 3 mice. Cell types are indicated and are represented by different colors.

### Flow cytometry

Flow cytometry was used to assess immune cell profiles in TG. Single cell suspensions were first stained for viability using Zombie NIR™ Fixable Viability Kit (BioLegend; San Diego, CA) for 20 min at room temperature in phosphate buffer solution pH 7.2 (PBS) combined with FcR blocking antibody (1 μg, clone 2.4G2, BioXCell; Lebanon, NH) to block non-specific binding. Cells then were washed with 2% FBS/PBS and stained with antibodies against surface antigens for 30 min on ice. Fluorochrome-conjugated antibodies against mouse CD45 (clone 30-F11), CD1-tet (1B1), CD3 (145-2C11), B220 (RA3-6B2), CD11b (M1/70), CD24 (M1/69), CD64 (X54-5/7.1), CD11c (N418), NK1.1 (PK136), TCRβ (H57-597), MHC-II (M5/114.15.2), Ly-6G (1A8), and Ly-6C (KH1.4) were purchased from BioLegend (San Diego, CA), eBioscience (San Diego, CA) or BD Biosciences (San Jose, CA). Flow cytometry was performed using Celesta or LSRII cytometer (BD Biosciences; San Jose, CA). Data were analyzed using FlowJo LLC v10.6.1 software.

Gating strategy to select immune populations in TG was modified from previously described approach (Yu et al., 2016) and shown in *Figure 6*. Briefly, live/singlets/CD45^+^ cells were gated using the markers listed below to define specific cell populations: monocytes (Mo, CD11b^+^/MHCII^lo^/SSC^lo^/CD64^+^); all macrophages (Mph, CD11b^+^/MHCII^hi^/CD64^+^/CD24^lo^/CD11c^-^); macrophage type-1 (Mph-1, Ly6C^+^/CCR2^+^); macrophage type-2 (Mph-2, Ly6C^+^/CCR2^+^); macrophage type-3 (Mph-3, Ly6C^-^/CCR2^-^/CX3CR1^+^/MHCII^+^); macrophage type-4 (Mph-4, Ly6C^-^/CCR2^-^/CX3CR1^+^/MHCII^-^); Ly6G^+^ neutrophils (Nph, CD11b^+^/Ly6G^+^/Ly6C^+^); dendritic cells (DCs; CD11^b+^/CD64^-^/CD24^hi^/MHCII^hi^/CD11c^+^); eosinophils (Eos; CD11b^+^/MHCII^hi^/CD64^-^/NK1.1^-^/CD3^-^); natural killer cells (NK; NK1.1^+^/TCRβ^-^); B cells (B, B220^+^/CD11^b-^/CD11^c-^); and T cells (T, CD3^+^/CD11b^-^/CD11c^-^).

### Immunohistochemistry (IHC)

For IHC, we used naïve wild-type, as well as Ccr2^RFP^Cx3cr1^GFP^ and Col1a2^cre^/Ai14^fl/-^ reporter male mice. TG tissues from 4% paraformaldehyde perfused mice were isolated, post-fixed with 4% paraformaldehyde for 1 h, cryoprotected with 10% and then 30% sucrose in phosphate buffer overnight, embedded in Neg 50 (Richard Allan Scientific, Kalamazoo, MI); and 25-30 μm cryo-sectioned. IHC was carried out as previously described (Lindquist et al., 2021). The following antibodies were used: anti-Iba1 rabbit polyclonal (Thermo Fisher Scientific; San Diego, CA; catalogue PA5-27436; 1:300); anti-s100a8 rat IgG2B monoclonal (R&D Systems; clone # 335806; Cat: MAB3059-SP 1:200) and anti-NFH chicken polyclonal (Novus Biological; Cat: NB300-217; 1:2000). Donkey Alexa Fluor secondary antibodies were from Jackson Immuno-Research (1:200; West Grove, PA). Control IHC was performed on tissue sections processed as described but either lacking primary antibodies or lacking primary and secondary antibodies. Images were acquired using a Keyence BZ-X810 All-in-One Fluorescent Microscope (Keyence, Itasca, IL). Gain setting was constant during acquisition, and it was established on no primary control slides. All Images taken were z-stack images and were processed with Adobe Photoshop CS2 software. Cells on micro-photography were counted within entire field captured with 20x objective.

### Statistical analysis

GraphPad Prism 8.0 (GraphPad, La Jolla, CA) was used for all non-single RNA-seq related statistical analyses of data. Data in the figures are mean ± standard error of the mean (SEM), with “n” referring to the number of animals per group. Differences between groups with one variable were assessed by chi-square analysis with Fisher’s exact test, unpaired t-test or regular 1-way ANOVA with Bonferroni’s post-hoc tests, each column was compared to all other columns. A difference was accepted as statistically significant when p<0.05. Interaction F ratios, and the associated p values are reported.

## Results

### Single-cell RNA sequencing of TG non-neuronal cells

We carried out two replicates for scRNA-seq on 12 left and right TG from 6 naïve, wild-type C57BL/6 male mice. Bilateral TG from 3 mice were combined per replicate. For first experiment, single-cell suspension preparation had Percoll step (see “Material and Method”). For second experiment, single-cell suspension preparation had two cycle FACS sorting step for isolation of viable cells (see “Material and Method”). For first replicate experiment, 2859 TG cells were present in scRNA-seq and an average of 1211 genes were detected per cell. For second replicate experiment, 6362 TG cells were present in scRNA-seq and an average of 1580 genes were detected per cell. All TG cells in each experiment were clustered together. We clustered and classified cell types using Loupe browser interface based tool from 10X and K-means for the first replicate and graph-based for the second replicate. Clusters enriched with mitochondrial gene were excluded. Clusters enriched for the expression of Pirt were assigned as sensory neuronal cluster(s), remaining clusters were classified as non-neuronal cells.

We detected 9 clusters (Cluster-1 has 471 cells, Cluster-2 351, Cluster-3 288, Cluster-4 239, Cluster-5 192, Cluster-6 168, Cluster-7 72, Cluster-8 37 and Cluster-9 21 cells) in the first experimental run/ replicate (*Fig 1A*), and 11 clusters (Cluster-1 has 1126 cells, Cluster-2 815, Cluster-3 618, Cluster-4 561, Cluster-5 446, Cluster-6 382, Cluster-7 356, Cluster-8 333, Cluster-9 311, Cluster-10 185 and Cluster-11 81 cells) in the second run/replicate (*Fig 1B*). Cell types were identified according to significant enrichment of markers (Padj<0.05; FC>5) and also by specificity in expression for certain genes (FC>10). It could be noted that neuronal cluster was not detected in the second experimental trial. This could be explained by no neuron Percoll enriching step in the second trial. This also indicate that scRNA-seq has substantial preference in sequencing non-neuronal cells over sensory neurons.

### Glial cells in TG

Glial cell sub-types were revealed after analysis of the first trial (*Fig 1A*), while second trial and analysis revealed only single glial group (*Fig 1B*). Schwann cells were recognized by an enrichment with *Mpz* (*Figs 1A, 2*; *Table 1*). SGC cells were distinguished by a specific marker - *Fabp7* (*Fig 1A; 2*; *Table 1*). *Fabp7* as SCG marker was noted in previous report as well (Chu et al., 2023). Suggested *Apoe* as a SGC marker was found to also be expressed at high levels in macrophages in our study (*Table 1*) (Yang et al., 2022;Chu et al., 2023). *Plp1* is also not suitable as a SGC marker, since it is present at an equal level in both Schwann cells and SGC (*Figs 1A, 2, Table 1*) (Kim et al., 2016;Kim et al., 2021). Interestingly, we could not distinguish between myelinating and non-myelinating Schwann cells using *Ncmap, Prx, Scn7a* and *Cdh19* (Yang et al., 2022;Chu et al., 2023). Thus, *Ncmap* and *Prx* were specific for Schwann cells and absent in SCG, *Cdh19* was only SCG cells and *Scn7a* was mainly in TG neurons (Supplementary Material). Despite glial cells contain distinct markers, it could be noted that low-to-moderate level of expression of these genes have been reported in sensory neurons (Usoskin et al., 2015;Sharma et al., 2020;Yang et al., 2022). Overall, our scRNA-seq data have shown that TG glial cells are divided into two groups: Schwann cells and SGC, which have distinct transcriptional profiles. Previous publications showed both Schwann cells and SGC could be clustered into variety of subtypes. Thus, Schwann cells were classified into myelinating and non-myelinating, while SGC types were divided into general resident, sensory, IEG and immune responsive groups (Mapps et al., 2022;Yang et al., 2022;Chu et al., 2023).

**Figure 2.**
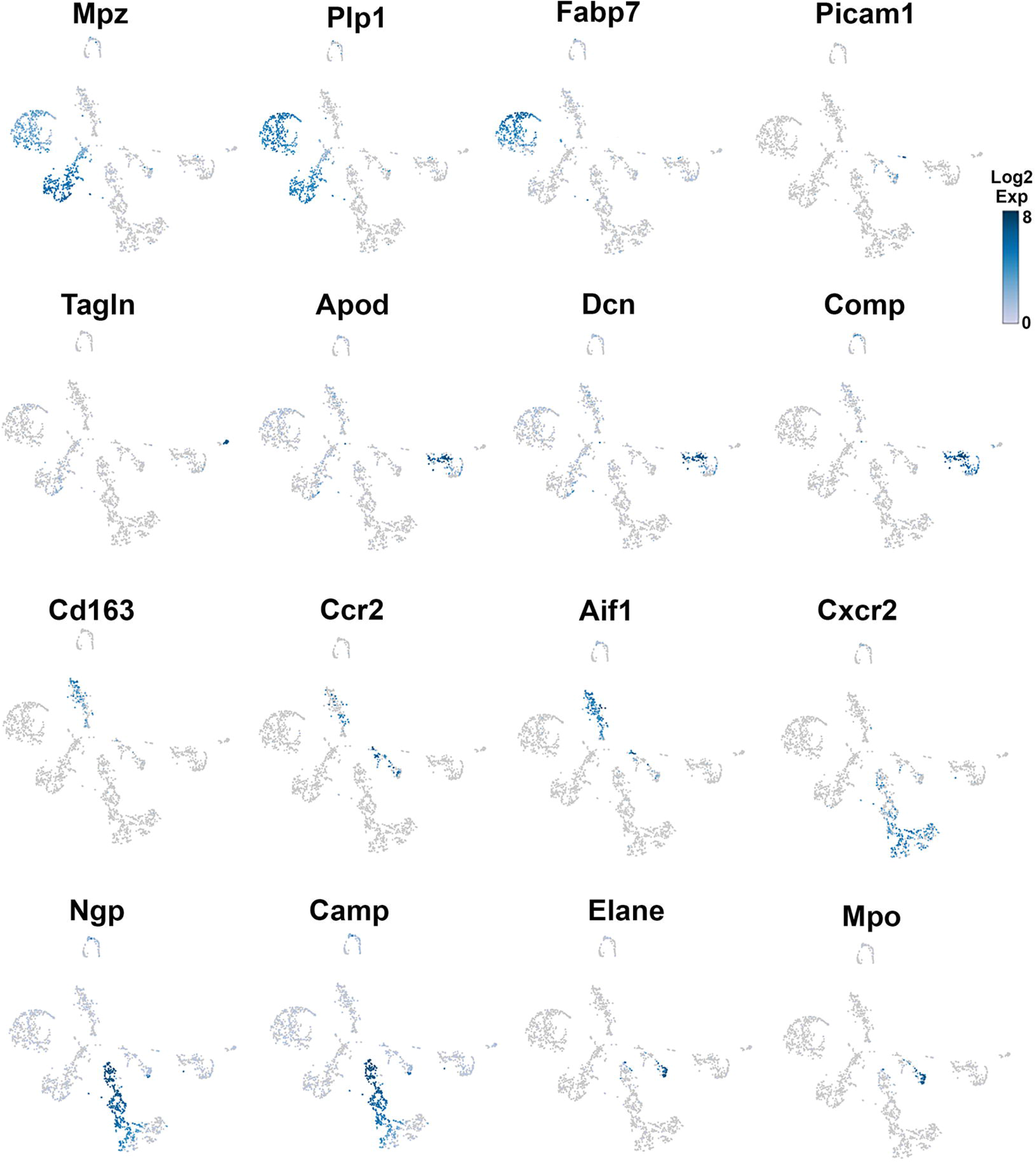
Expression of markers in TG non-neuronal cells (Experiment-1) The expressions of select cell-type marker genes for TG non-neuronal cells clustered in the experiment-1 are displayed in individual cells and within clusters. Marker genes are indicated.

**Table 1.** Markers for non-neuronal cells in TG.

| ID | SchC | SGC | Mus | Endo | Fibro1 | Fibro2 | Mph1 | Mph2 | Mph3 | Neu1 | Neu2 | Neu3 | Neu4 |
| --- | --- | --- | --- | --- | --- | --- | --- | --- | --- | --- | --- | --- | --- |
| Mpz | 152 | 8 | - | - | - | - | - | - | - | - | - | - | - |
| Mbp | 27 | 1.8 | - | - | - | - | - | - | - | - | - | - | - |
| Plp1 | 13 | 15 | - | - | - | - | - | - | - | - | - | - | - |
| Apoe | 1.6 | 188 | 1.6 | - | 27 | - | 274 | 207 | 217 | - | - | - | - |
| Fabp7 | - | 26 | - | - | - | - | - | - | - | - | - | - | - |
| Tie | - | - | - | 1 | - | - | - | - | - | - | - | - | - |
| Pecam1 | - | - | - | 2.7 | - | - | - | - | - | - | - | - | - |
| Tagln | - | - | 46 | - | - | - | - | - | - | - | - | - | - |
| Tpm1 | - | - | 18 | - | - | - | - | - | - | - | - | - | - |
| Myh11 | - | - | 17 | - | - | - | - | - | - | - | - | - | - |
| Col1a2 | - | - | - | - | 7.7 | 13 | - | - | - | - | - | - | - |
| Apod | - | - | - | - | 293 | 9 | - | - | 30 | - | - | - | - |
| Dcn | - | - | - | - | 174 | 22 | - | - | 9 | - | - | - | - |
| Vtn | - | - | - | - | 3.6 | - | - | - | - | - | - | - | - |
| Ccl11 | - | - | - | - | 7.8 | - | - | - | - | - | - | - | - |
| Mgp | - | - | 8.5 | 8.6 | 16 | 117 | - | - | 6 | - | - | - | - |
| Comp | - | - | - | - | - | 5.3 | - | - | - | - | - | - | - |
| IL33 | - | - | - | - | 2.7 | 0.5 | - | - | - | - | - | - | - |
| Pdgfra | - | - | - | - | 1.1 | - | - | - | - | - | - | - | - |
| Pdgfrb | - | - | - | - | 1.1 | - | - | - | - | - | - | - | - |
| Csf1r | - | - | - | - | - | - | 6 | 4 | 4 | - | - | - | - |
| Ccr2 | - | - | - | - | - | - | - | 1.2 | - | - | - | - | 2.1 |
| Cx3cr1 | - | - | - | - | - | - | 3.5 | 2.5 | 0.8 | - | - | - | - |
| Cd68 | - | - | - | - | - | - | 2.5 | 2.3 | 2.1 | - | - | - | - |
| Cd163 | - | - | - | - | - | - | 1.8 | 0.5 | 0.8 | - | - | - | - |
| Aif | - | - | - | - | - | - | 3.3 | 3.6 | 6 | - | - | - | - |
| Cxcl9 | - | - | - | - | - | - | - | - | 1.4 | - | - | - | - |
| Csf3r | - | - | - | - | - | - | - | - | - | 4.3 | - | 0.8 | - |
| Cxcr2 | - | - | - | - | - | - | - | - | - | 2.8 | - | 1.5 | - |
| Cxcr4 | - | - | - | - | - | - | - | - | - | 2.6 | - | 0.9 | - |
| Cxcl2 | - | - | - | - | - | - | 2.3 | 10 | - | 36 | - | 4 | 3 |
| S100a8 | - | - | - | - | - | - | - | - | - | 319 | 590 | 615 | 16 |
| S100a9 | - | - | - | - | - | - | - | - | - | 462 | 858 | 914 | 20 |
| Ngp | - | - | - | - | 1.3 | - | - | - | - | - | 240 | 99 | 2.3 |
| Ly6G | - | - | - | - | - | - | - | 0.9 | - | - | 4.8 | 4 | - |
| Cd177 | - | - | - | - | - | - | - | - | - | - | 2.7 | 2.7 | - |
| Camp | - | - | - | - | - | - | - | - | - | - | 204 | 56 | 3 |
| Ltf | - | - | - | - | - | - | - | - | - | - | 40 | 19 | - |
| Elane | - | - | - | - | - | - | - | - | - | - | 2 | - | 5.5 |
| Mpo | - | - | - | - | - | - | - | - | - | - | - | - | 2.6 |
| Pycard | - | - | - | - | - | - | - | - | - | - | - | - | 1.25 |
| Ramp1 | - | - | 3 | - | - | - | - | - | - | - | - | - | 1.1 |
| Ccl5 | - | - | - | - | - | - | - | - | - | - | - | - | 3 |
| Cd74 | - | - | - | - | 1.4 | 1.8 | 90 | 118 | - | 59 | 19 | - | - |
| Ccl2 | - | - | - | - | 2.8 | - | 2.5 | 3.2 | - | - | - | - | - |
| Itgam | - | - | - | - | - | - | 0.6 | 0.4 | 0.35 | 1.2 | 1.3 | 1.7 | 0.5 |
| Ptpcr | - | - | - | - | - | - | 1.6 | 1.4 | 0.6 | 5.9 | 1.3 | 2.7 | 2.4 |
| Lyz2 | - | - | - | - | - | - | 66 | 69 | 54 | 30 | 77 | 56 | 63 |
| Itgax | - | - | - | - | - | - | - | - | - | - | - | - | - |
| Cd3d | - | - | - | - | - | - | - | - | - | - | - | - | - |
| Cd22 | - | - | - | - | - | - | - | - | - | - | - | - | - |
| Il2rb | - | - | - | - | - | - | - | - | - | - | - | - | - |

- shows expression *less than 0.5*; **SchC** is Schwann cells; **SGC** is satellite glial cells; **Mus** is smooth muscle. **Endo** is endothelial cells; **Fibro1** is Apod<sup>+</sup>/Dcn<sup>high</sup> fibroblasts; **Fibro2** is Comp<sup>+</sup>/Mgp<sup>high</sup> fibroblasts; **Mph1** is CD163<sup>high</sup> macrophages (Mph); **Mph2** is Cx3cr1<sup>+</sup> Mph; **Mph3** is Aif1<sup>high</sup> Mph; **Neu1** is Cxcr2<sup>high</sup>/Cxcr4<sup>+</sup> neutrophils (Neu); **Neu2** is Ly6G<sup>+</sup> Neu-2; **Neu3** is Ly6G<sup>+</sup> Neu-2; **Neu4** – Mpo<sup>+</sup>/Elane<sup>high</sup>/Ly6G<sup>-</sup> Neu.

### Fibroblasts, smooth muscle and endothelial cells in TG

Different types of fibroblasts were categorized after analysis of second (*Fig 1B*), but not in the first trial (*Fig 1A*). Fibroblasts identified as Col1a2^+^ cells were represented by two distinct clusters (*Fig 1B*). The larger fibroblast subgroup, labeled as Fibro (Dcn), was *Apod^+^/Dcn^high^* and was distinguished by a combination of an enriched marker *Dcn* and a specific marker *Apod* (*Figs 1B, 3*; *Table 1*). Unlike *Dcn*, *Apod* is expressed by many sensory neurons (Usoskin et al., 2015;Sharma et al., 2020;Yang et al., 2022). The second subgroup, labeled as Fibro (Comp), was *Comp^+^/Mgp^high^*, was enriched with *Mgp* and contained specific marker *Comp* (*Figs 1B, 3; Table 1*), and was previously reported as myofibroblasts (Chu et al., 2023).

**Figure 3.**
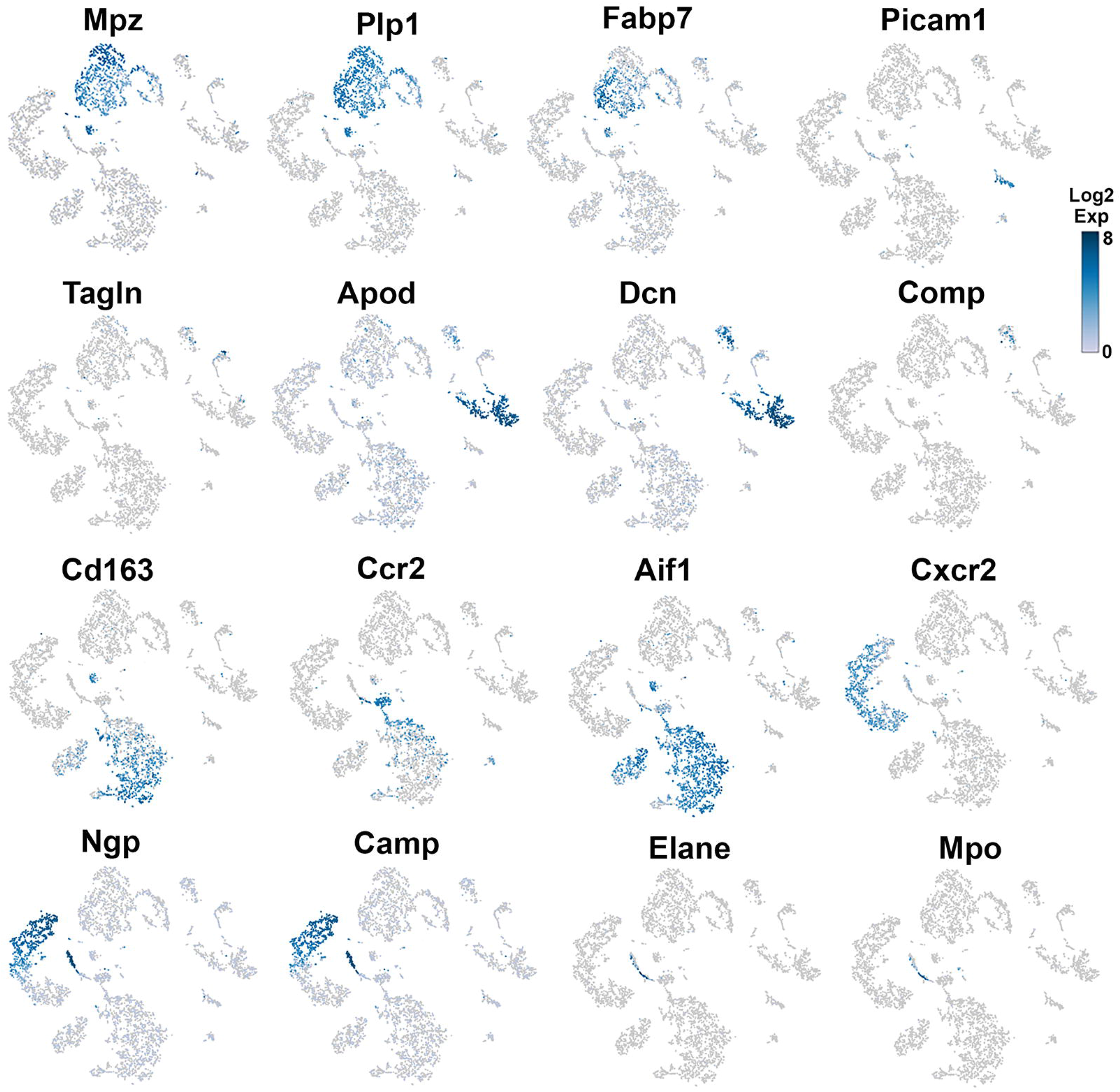
Expression of markers in TG non-neuronal cells (Experiment-2) The expressions of select cell-type marker genes for TG non-neuronal cells clustered in the experiment-2 are displayed in individual cells and within clusters. Marker genes are indicated.

We and others also identified small number of cells representing either smooth muscle or endothelial vascular cells (*Figs 1A, 1B*) (Yang et al., 2022;Chu et al., 2023). Smooth muscle vascular cells contained a specific marker *Tagln*; and endothelial vascular cells defined by standard *Pecam1/CD31* marker (*Figs 1B, 1A, 2, 3*; *Table 1*). In summary, there is an agreement that TG have two types of fibroblasts (Apod^+^/Dcn^high^ and Comp^+^/Mgp^high^) as well as vascular smooth muscle and endothelial cells (Sharma et al., 2020;Yang et al., 2022;Chu et al., 2023).

### Immune cells in TG

Immune cells in TG were reported as one single group (Sharma et al., 2020;Yang et al., 2022;Chu et al., 2023), though DRG immune cells were split into three groups: macrophages (Mph), neutrophils (Neu) and B-cells (Renthal et al., 2020). Our first trial revealed a group of Mph and two groups of Neu (*Fig 1A*). Second trial, which did not enrichment for sensory neurons, contained more cells and deeper reading, split these groups into 3 Mph and 4 Neu groups (*Fig 1B*). Mph were recognized by expression of CD64 (aka *Fcgr1*). First group of Mph was enriched with *Cd163*, a marker for M2-type of Mph (Landis et al., 2018;Cutolo et al., 2022), but contained other M1 Mph markers, such as *Csfr1*, *Cx3cr1*, *Cxcr1* and *Cd68* (*Figs 1B, 3, Table 1*). Second group of Mph had specific marker, *Ccr2*, and was expressing *Csfr1*, *Cxcr1* and *Cd68* as well (*Figs 1B, 3, Table 1*). Third group of Mph could be classified as *Aif1*^high^ (aka Iba1), which also specifically expressed *Cxcl9* (*Figs 1B, 3, Table 1*). Third Mph group had low level of *Cd163* and no *Ccr2*, and expressed several fibroblast markers (*Fig 3, Table 1*).

Neu were recognized by expression of s100a8 and s100a9 (Foell et al., 2009;Di Ceglie et al., 2018). Four groups of Neu could be split into two domains Ly6G^-^ and Ly6G^+^ (*Table 1*). Ly6G^+^ Neu groups (Neu-2 and Neu-3) were similar and have had high level of *Cd177, Camp* and *Ngp* expressions (*Table 1*). Nevertheless, Neu2 and Neu-3 could be differentiated by transcriptional profiles and expressions of *Elane* in Neu-2 and *Cxcl2* and *Cxcr2* in Neu-3 (*Fig 1B, Table 1*). Ly6G^-^ Neu groups (Neu-1 and Neu-4) were substantially different from Ly6G^+^ Neu, and were dissimilar to each other (*Fig 1B, Table 1*). Thus, Neu-1 was enriched with *Cxcr4* and *Csf3r*, while Neu-4 was dominated with *Ccl5*, *Elane*, *Pycard* and *Mpo* (*Table 1*).

Next, we looked at expressions of standard immune cell markers in clusters of TG non-neuronal cells. A pan-immune cell marker, *Ptprc* (aka CD45), was expressed by all TG immune cells. However, the expression level was low (*Table 1*). A pan-myeloid cell marker *Itgam* (aka CD11b) was present at a very low levels in Mph and at low-to-moderate levels in Neu (*Table 1*). Mph and monocyte marker, *Ccl2*, was expressed by a subset of TG Mph and *Apod^+^/Dcn^high^* fibroblasts (*Table 1*). A myeloid cell marker, *CD74*, was subsets of TG Mph and Neu, and at lower levels in TG fibroblasts (*Table 1*). We found that most suitable and highly expressed marker for all TG immune cells was *Lyz2* (aka LyzM; *Table 1*). scRNA-seq showed that markers for other immune cell types, including *Itgax* (aka CD11c) for dendritic cells, *Il2rb* (aka CD122) for natural killer cells, *Cd3d* and *Cd3e* for T-cells and *Cd22* for B-cells are not present in TG (*Table 1*). Altogether, TG had exclusively a subset of myeloid cells, which could be sub-grouped into several Mph and Neu types with distinct transcriptional profiles.

### Visualizations of non-neuronal cells in TG by immunohistochemistry (IHC)

Glial cells were thoroughly visualized and quantified in sensory ganglia in many reports (Kim et al., 2016;Mapps et al., 2022). Mph and Neu were mainly visualized in DRG and were often presented as a single group (Latremoliere et al., 2015;Guan et al., 2016;Woolf et al., 2021;Yu et al., 2021;Caxaria et al., 2023). Here we investigated whether several groups of Mph, Neu and fibroblasts could be detected in TG. To do so, we have used two approaches: immunohistochemistry (IHC) and flow cytometry.

Based on our scRNA-seq data, different types of Mph could be distinguished by labeling TG from Ccr2^RFP^/Cx3cr1^GFP^ reporter with Iba1 antibodies. In congruence with scRNA-seq data, three subsets of Mph – Cx3cr1^+^/Ccr2^-^/Iba1^+^ (Mph-1), Cx3cr1^+^/Ccr2^+^/Iba1^+^ (Mph-2), and Cx3cr1^-^/Ccr2^-^/Iba1^+^ (Mph-3) - could be distinguished (*Figs 4A-A”’*, *Table 1*). scRNA-seq revealed 633 and 618 cells in the Mph-1 and Mph-2 clusters, respectively, and 356 cells in the Mph-3 cluster (*Fig 1B*). Cell counting showed that there were substantially lesser Ccr2^+^ than Cx3cr1^+^ and Iba1^+^ cells in TG (1-way ANOVA; F (4, 10) = 42.48; P<0.0001; n=3; *Fig 5*).

**Figure 4.**
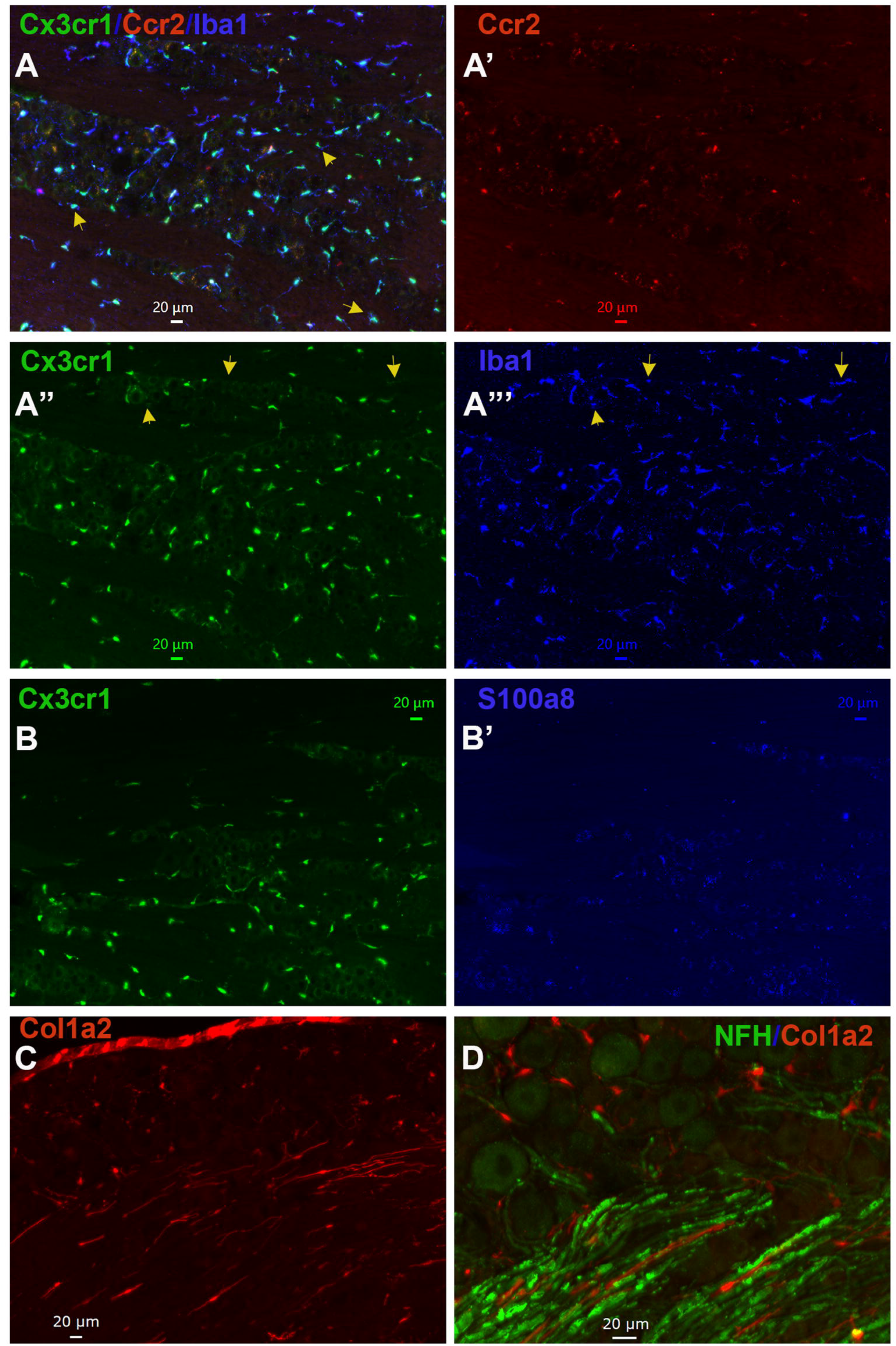
Representations of non-neuronal cells in TG. (**A-A”’**) Representative micro-photographs of Ccr2^RFP^/Cx3cr1^GFP^ reporter mouse TG sections show relative expressions of macrophage markers Cx3cr1 (green), Ccr2 (red) and Iba1 (blue). Yellow arrows on *the panels A* show Cx3cr1^+^/Iba1^+^/Ccr2^-^ macrophages in TG. Yellow arrows on *the panels A” and A”’* show Cx3cr1^-^/Iba1^+^/Ccr2^-^ macrophages in TG. (**B-B’**) Representative micro-photographs of Ccr2^RFP^/Cx3cr1^GFP^ reporter mouse TG sections show relative expressions of a macrophage marker Cx3cr1 and a neutrophil marker S100a8. (**C**) A representative micro-photograph of a Col1a2^cre^/Ai14^fl/^ reporter mouse TG sections shows expression of a fibroblast marker Col1a2. (**D**) A representative micro-photograph of a Col1a2^cre^/Ai14^fl/^ reporter mouse TG section shows relative expressions of a fibroblast marker Col1a2 and a A-fiber neuronal marker NFH. Scales are presented in each microphotograph.

**Figure 5:**
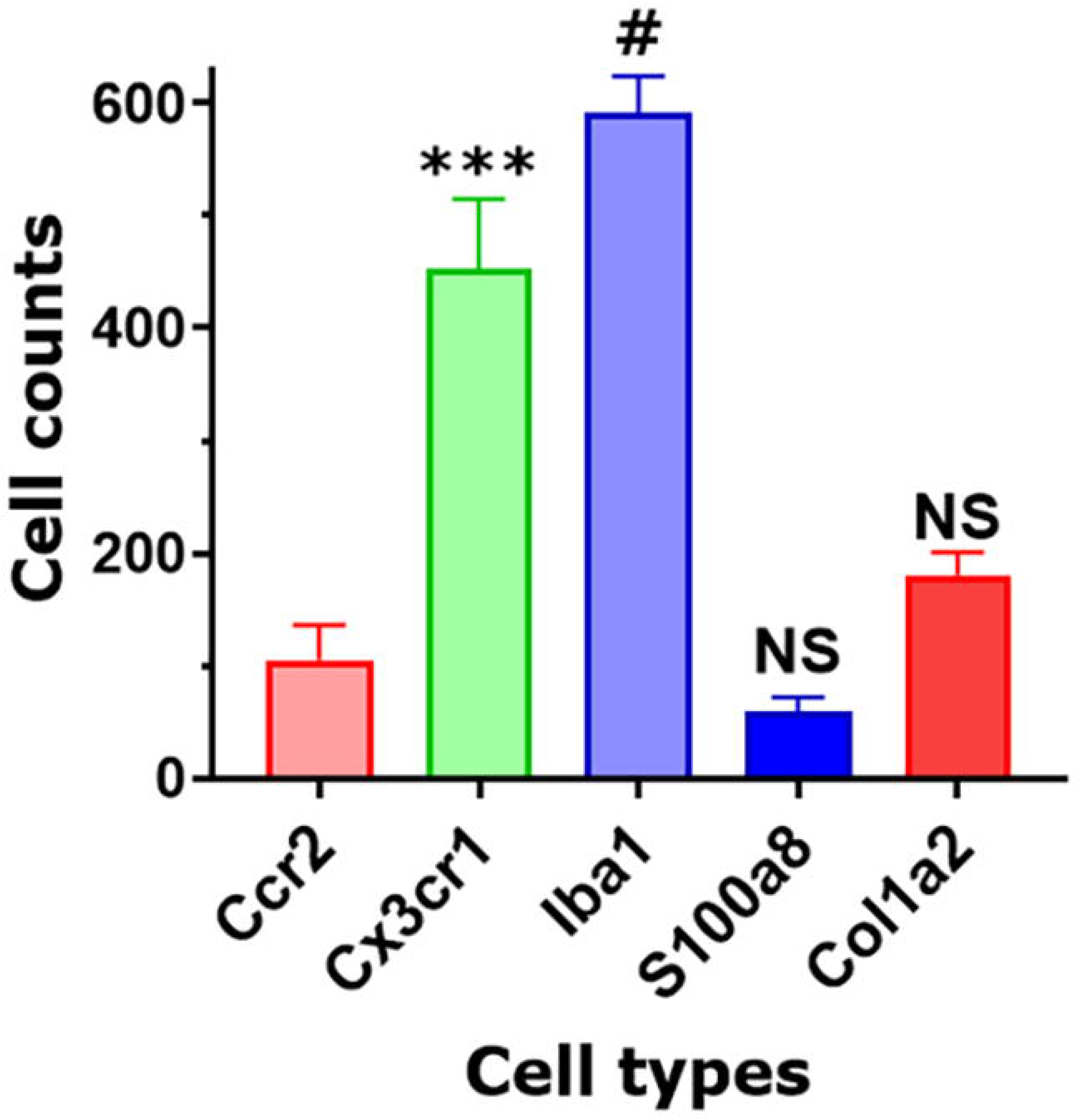
Non-neuronal cell count in TG sections. Counts of different types of non-neuronal cells in TG sections from Col1a2^cre^/Ai14^fl/^ and Ccr2^RFP^/Cx3cr1^GFP^ reporter mice. Every cell within a field of microscope equipped with x20 objective was counted. Statistic is 1-way ANOVA relatively to the Ccr2 column (NS p>0.05; *** p<0.001; # p<0.0001; n=3).

Highly expressing Neu marker s100a8 (*Table 1*) was detected in few TG cells (*Figs 4B, 4B’, 5*). This is surprising data considering at least similar numbers of Mph and Neu in TG were revealed by scRNA-seq (*Figs 1A, 1B*). We hypothesized that in naïve wild-type mice, Neu is mainly located in dura sheath covering entire TG (see the next section) (Caxaria et al., 2023).

Fibroblasts were visualized by IHC on TG from Col1a2^cre^/Ai14^fl/-^ reporter mice. Fibroblasts with distinct shapes compared to immune and glial cells were noted among neuronal cell bodies and myelinated nerve fibers in TG (*Figs 4C, 4D*). In summary, IHC can distinguish different types of TG Mph and visualize fibroblasts. IHC data also implied that Neu could be located in dura sheath covering TG.

### Detection of TG immune cells by flow cytometry

To further validate findings of scRNA-seq on immune cells, we have performed flow cytometry on single-cell suspensions generated from male mouse TG. TG was isolated without or with surrounding dura (see “Material and Methods”). DRG was also isolated from some animals. Overall flow cytometry gating strategy is presented in *Figure 6*. TG and DRG without dura are dominated by Mph (*Figs 7A, 7B*). Neu and T-cell levels are low in naïve wild-type mice (*Figs 7A, 7B*). Based on scRNA-seq and IHC results, we have design gating strategy to detect different types of Mph (*Fig 7C*). Nether TG nor DRG had Ccr2^+^/Ly6C^+^/MHCII^+^/Cx3cr1^+^ Mph, which could be classified as inflammatory macrophages (*Figs 7D, 7E*). TG and DRG also lacked Mph without Ly6C and Cx3cr1 (Ccr2^+^/Ly6C^-^/MHCII^+^/Cx3cr1^-^; *Figs 7D, 7E*). Both TG and DRG had Mph lacking Ly6C or Ly6C and Ccr2 markers: Ccr2^+^/Ly6C^-^/MHCII^+^/Cx3cr1^+^ and Ccr2^-^/Ly6C^-^/MHCII^+^/Cx3cr1^+^, respectively (*Figs 7D, 7E*). Finally, Mph with only Cx3cr1 (Ccr2^-^/Ly6C^-^/MHCII^-^/Cx3cr1^+^) were present in TG, but not DRG (*Figs 7D, 7E*).

**Figure 6:**
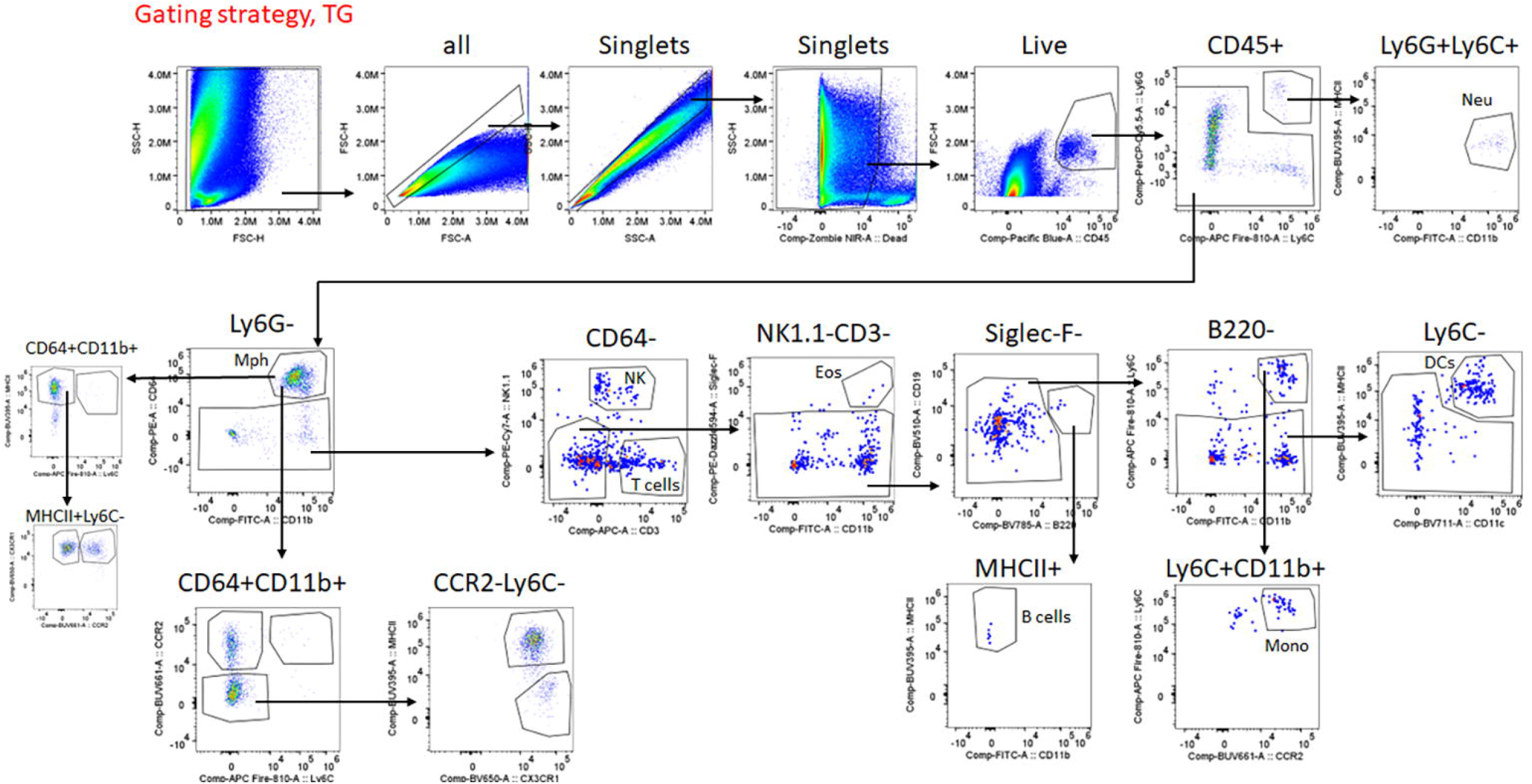
Gating strategy for flow cytometry. Sequence of gating events leads to selection of live singlets, and then separation of CD45^+^ cells. CD45+ cells, specific markers and gating strategy are used to calculate numbers of different immune cells. Immune cell types and markers are indicated. Sequence of gaining events are marked by arrows.

**Figure 7:**
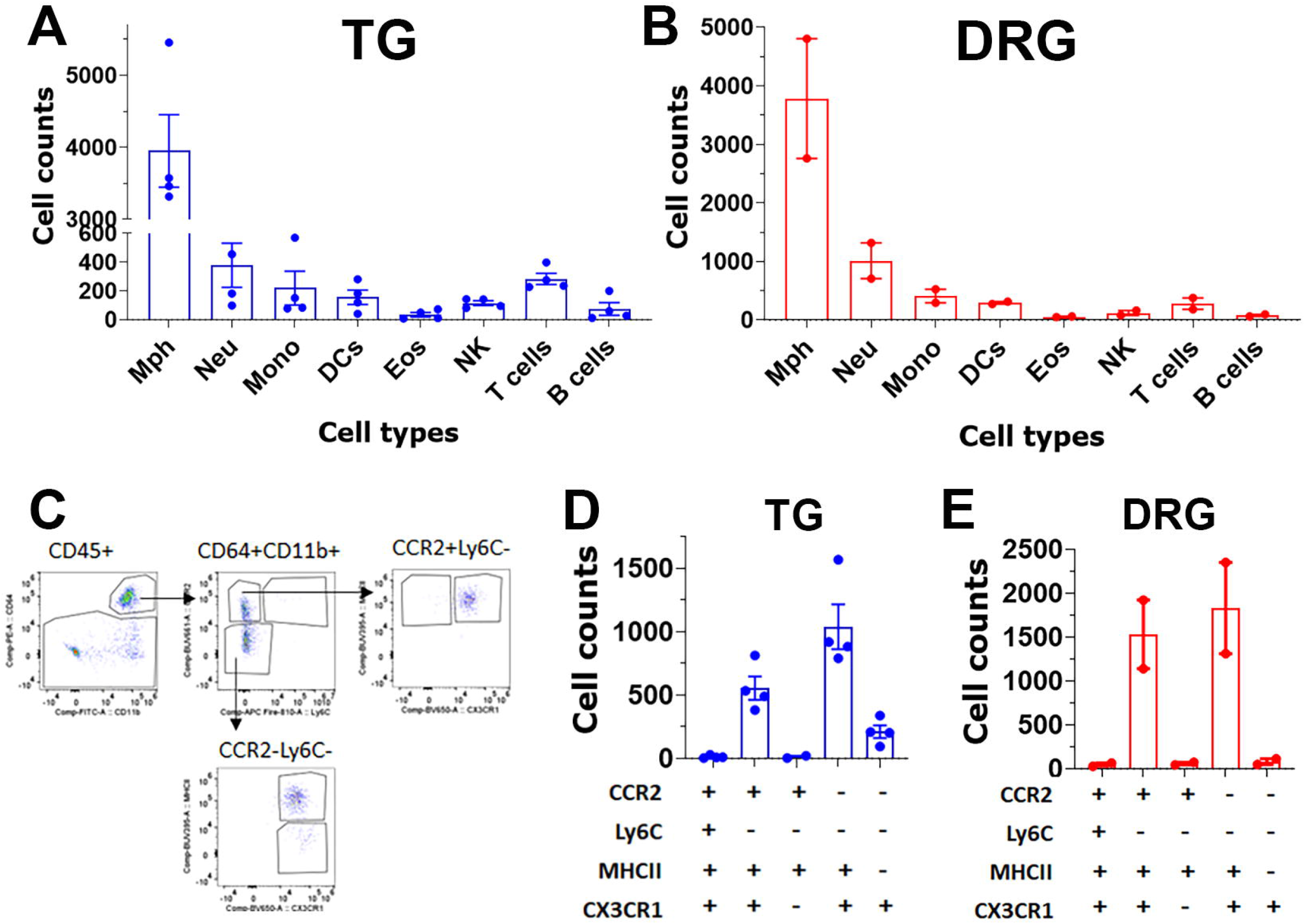
Immune cell profiles in TG and DRG isolations without surrounding dura. Normalized (by live cell numbers) immune cell counts in the TG (the panel **A**) and DRG (the panel **B**) without surrounding dura isolated from naïve WT male mice. Mph – macrophages; Neu – neutrophils; Mono – monocytes; DCs – dendritic cells; Eos – eosinophils; NK – natural killer cells; B-cells; and T-cells (n=3-4). (**C**) Gating strategy to isolate variety types of macrophages. Normalized (by live cell numbers) cell counts for different types of macrophages in TG (panel **D**) and DRG (panel **E**) without surrounding dura. Used markers are indicated beneath panel D and E.

Isolation of TG with surrounding dura, which is described in the “Material and Method” section, dramatically changed immune cell profiles (*Fig 8A*). Neu dominated CD45^+^ cells (*Fig 8A*). Besides Neu, isolation of TG with dura led to an increase in other immune cells, especially monocytes (MO; *Fig 8A*). Interestingly, isolation TG with dura did not alter ganglion Mph and dendritic cells (DCs) profiles (*Figs 8B, 8C* vs *Figs 7D, 7E*). Thus, TG with dura had Ccr2^+^/Ly6C^-^/MHCII^+^/Cx3cr1^+^ and Ccr2^-^/Ly6C^-^/MHCII^+^/Cx3cr1^+^ Mph groups (*Figs 8C, 8D*). However, it appears that Mph with only Cx3cr1 (Ccr2^-^/Ly6C^-^/MHCII^-^/Cx3cr1^+^) were more present in TG with dura compare with TG without dura (*Figs 8C vs 7E*). Overall, flow cytometry study validated scRNA-seq data on diversity of Mph and Neu in TG, and showed that Neu is mainly present in dura surrounding TG from naïve WT mice.

**Figure 8:**
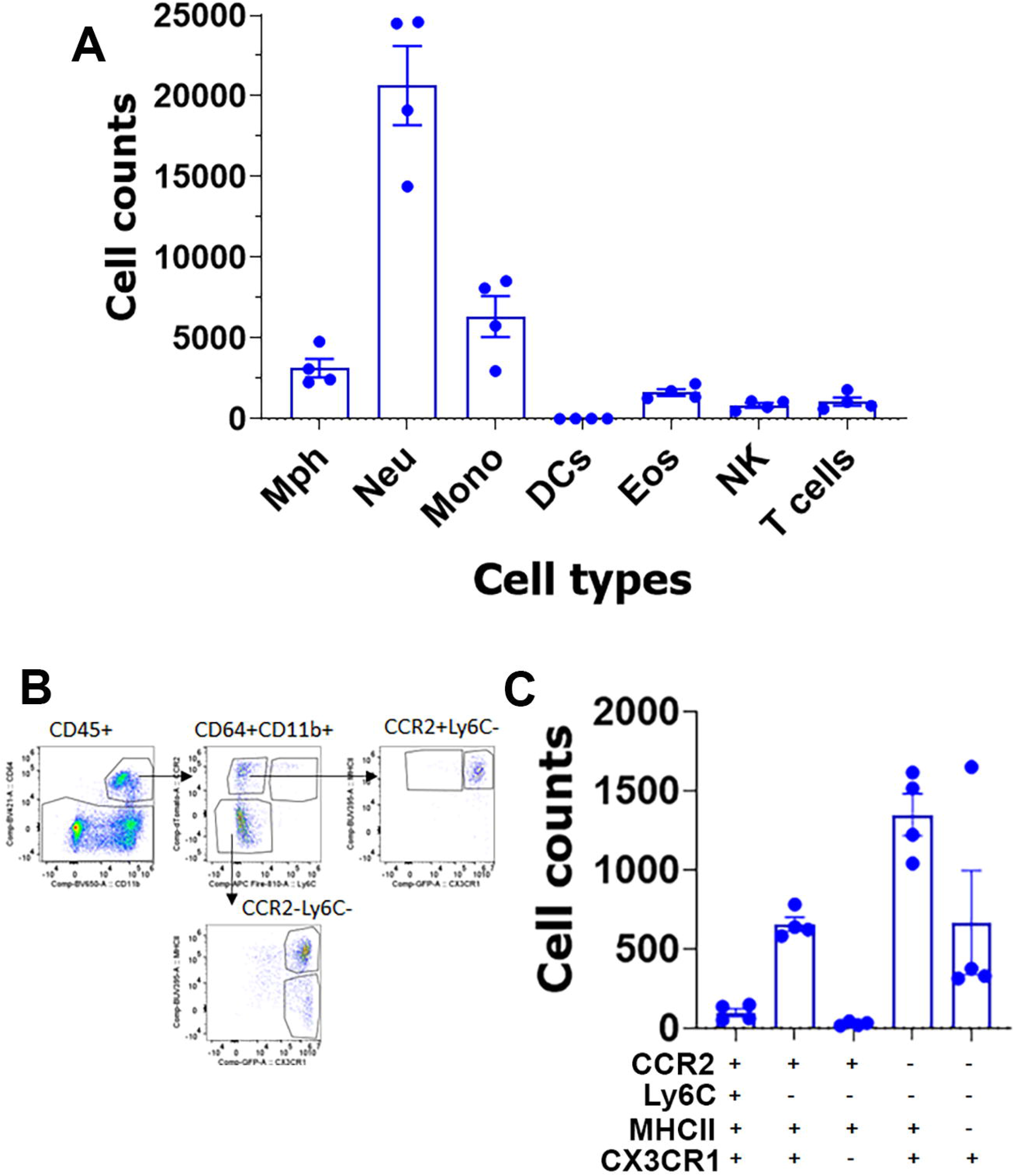
Immune cell profiles in TG isolations with surrounding dura. (**A**) Normalized (by live cell numbers) immune cell counts in the TG with surrounding dura isolated from naïve Ccr2^RFP^/Cx3cr1^GFP^ reporter male mice. Mph – macrophages; Neu – neutrophils; Mono – monocytes; DCs – dendritic cells; Eos – eosinophils; NK – natural killer cells; and T-cells (n=3-4). (**B**) Gating strategy to isolate variety types of macrophages. (**C**) Normalized (by live cell numbers) cell counts for different types of macrophages in TG with surrounding dura. Used markers are indicated below X-axis.

## Discussion

Plasticity of non-neuronal cells in TG and DRG during a variety of acute and chronic pain conditions were reported in multiple publications. This plasticity of glial (Huang et al., 2013;Donnelly et al., 2020;Gazerani, 2021;McGinnis and Ji, 2023) and immune cells (Ji et al., 2016;Yu et al., 2020;Domoto et al., 2021;Lesnak et al., 2023) has essential contributions to progression of pain in many diseased states. It is presumed that transcriptional changes in sensory ganglion non-neuronal cells could eventually lead to production of plethora of mediators. It has been shown that these mediators can sensitize sensory neurons (Ji et al., 2016;Donnelly et al., 2020;Gazerani, 2021) by direct communication with sensory neurons (Kim et al., 2016) and/or by regulating a plethora of neuronal channels, which could result in changing neuronal excitability and gating properties (Domoto et al., 2021;Haberberger et al., 2023;McGinnis and Ji, 2023).

This essential role of non-neuronal ganglion cells in the regulation of sensory neuronal excitability and the development of pain conditions elevated information on transcriptional profiles for non-neuronal sensory ganglion cells to a critically important level. Thus, this data could be used for calculating interactomic network between sensory neurons and non-neuronal cells (Wangzhou et al., 2021). This dataset could also be baseline in investigation of TG non-neuronal cell plasticity in different pain models for head and neck area. RNA-seq on single cell level was employed in several publications to gain this important information and knowledge on transcriptional profiling of non-neuronal cells in mouse TG (Sharma et al., 2020;Mapps et al., 2022;Yang et al., 2022;Chu et al., 2023). Results of these independent studies have substantial overlap as well as differences. These differences are unavoidable and could be dictated by nuclei/cell isolation approaches, numbers of sequenced cells, sequencing depth and clustering analysis (Drokhlyansky et al., 2020;Renthal et al., 2020;Sharma et al., 2020;Mapps et al., 2022;Yang et al., 2022). Generally, multiple independent repetitions of scRNA-seq positively contributes to generation of accurate single-cell profiles for tissues at naïve and pathological conditions. Accordingly, we performed two different non-neuronal cell isolation and two separate replicates (reported separately in *Figs 1-3*) of scRNA-seq to generate transcriptional profiles for non-neuronal TG cells. Then, we integrated previously published and our data (*Fig 9*).

**Figure 9.**
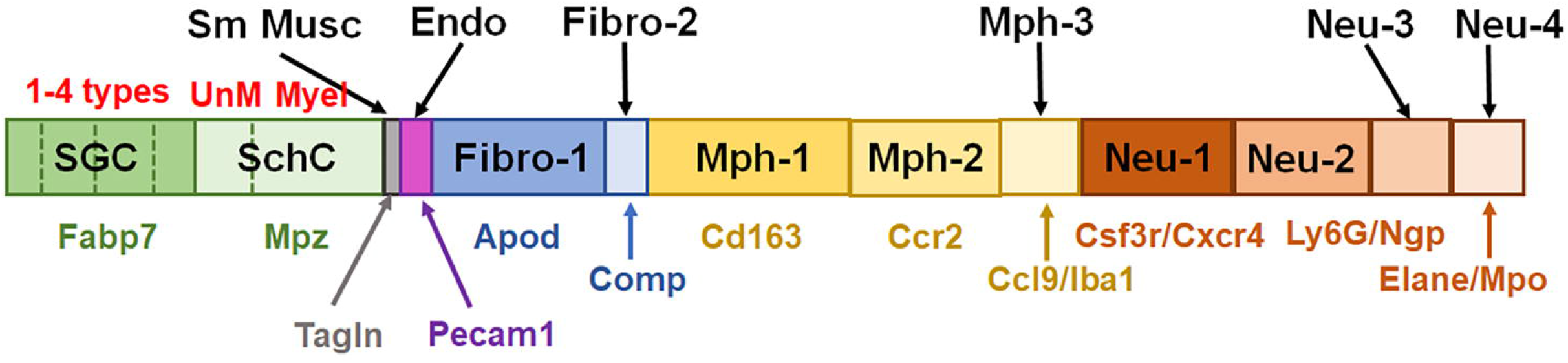
Schematic for representation of non-neuronal cells in TG. This schematic summarize published and generated here data. SGC – satellite glial cells are represented by 4 different subtypes. SchC – Schwann cells are represented by two subtypes: unmyelinated (UnM) and myelinated (Myel). Sm Musc – vascular smooth muscle cells. Endo - vascular endothelial cells. Fibro – fibroblasts are divided on two sub-groups: Fibro-1 and Fibro-2. Mph – macrophages are divided on three sub-groups: Mph-1, Mph-2 and Mph-3. Neu – neutrophils are divided on four sub-groups: Neu-1, Neu-2, Neu-3 and Neu-4. Gene markers for each TG non-neuronal cell type is indicated underneath of bars with the same color theme as bars.

There are several types of glial cells in TG (*Fig 1*). Schwann cells could be assigned to one of two categories: myelinated and non-myelinated (Yang et al., 2022;Chu et al., 2023). SCG could also be divided on four subsets: general resident, sensory, IEG and immune responsive (Mapps et al., 2022). Specific marker for all SGC sub-groups is *Fabp7* and for all Schwann cell sub-groups is Mpz. Suggested *Apoe* as a SGC marker is expressed at high levels in macrophages (*Table 1*) (Yang et al., 2022;Chu et al., 2023). *Plp1* is also not suitable as a SGC marker, since it is present at an equal level in Schwann cells and SGC (*Table 1*) (Kim et al., 2016;Kim et al., 2021). Moreover, low-to-moderate level of expression of glial markers have been reported in sensory neurons (Usoskin et al., 2015;Sharma et al., 2020;Yang et al., 2022). Overall, we could not find viable strategy in controlling specific types of glial cells without affecting other cell types, including sensory neurons.

Among TG non-neuronal cells are small numbers of vascular smooth muscle and endothelial cells (*Fig 1*). Two types of fibroblasts were reported previously (Yang et al., 2022;Chu et al., 2023). Our scRNA-seq data concur with this finding (*Fig 1*). We could suggest *Apod* as a gene marker for one group of fibroblasts and *Comp* for another group (*Fig 1*, Table 1). *Dcn* could be another good choice for a Fibro-1 group marker, since *Apod* is expressed on substantial levels in sensory neurons (Usoskin et al., 2015;Sharma et al., 2020;Yang et al., 2022). However, *Mgp* as a proposed marker for Fibro-2 is present on significant levels in other TG non-neuronal cells (*Table 1*). Additionally, it appears that most appropriate marker to manipulate all fibroblasts is Col1a2. Our IHC data have also shown that fibroblasts are among sensory neuronal cell bodies in TG as well as within bunches of nerve fibers.

Importance of sensory ganglion immune cells in the regulation of the development of pain conditions is well documented in many reports (see “Introduction”). ScRNA and snRNA-seq showed only one immune cell cluster in TG (Sharma et al., 2020;Yang et al., 2022;Chu et al., 2023). Interestingly, DRG contained macrophage, neutrophil and B-cell clusters (Renthal et al., 2020). Our data indicate that there are at least 7 immune cell clusters in TG (*Fig 12, Table 1*). IHC and flow cytometry confirmed presence of three type of residential macrophages (Cd64^+^) in TG, which could be recognized by Cd163, Ccr2 and Iba1 markers (*Figs 3, 4, Table 1*), and separated by flow cytometry (*Figs 7E, 8C*). Interestingly, Cd163 is a marker for M2 macrophages. Moreover, Cd206/Mrc1 and Il10rb as two recognized M2 markers are also expressed on Mph1 (Supplementary Material). scRNA-seq data clustering also revealed 4 types of neutrophils (s100a8^+^) which are either Ly6G^+^ or Ly6G^-^. Surprisingly, s100a8 antibodies, which labeled neutrophils in many tissues, did not show strong signal within naïve mouse TG. Neutrophils were mainly located in perineuronal/dura sheath surrounding DRG, but infiltrated DRG during myalgia (Caxaria et al., 2023). Similarly, flow cytometry revealed plenty of neutrophils in TG isolation containing surrounding dura (*Fig 8A*). Ly6G^+^ neutrophils were clustered into two groups, but we could not find specific markers distinguishing Neu-2 from Neu-3 sub-types (*Table 1*). Ly6G^-^ neutrophils could be readily differentiated into two groups using *Csf3r* or *Cxcr4* for Neu-1 and *Elane*, *Mpo* or *Pycard* for Neu-4 (*Table 1*).

In conclusion, comprehensive transcriptional profile data of TG non-neuronal cells generated by our and several independent studies establish foundation for detailed studies on regulation of a variety pain conditions by these cell sub-types. Detailed transcriptional profiles for each group of cells and independent replicates generated in this study are provided as a Supplementary Material. Thus, outlined here information could allow generating novel tools (mouse lines) for selective manipulations of particular sub-types on TG non-neuronal cells and evaluate cell plasticity during pain conditions. This in turn could address understudies areas of pain research by building the molecular basis for mechanisms controlling chronicity of a variety pain conditions for head and neck area.

## Abbreviations

TG: trigeminal ganglia
scRNA-seq: single-cell
RNA: sequencing
snRNA-seq: single-nuclear
RNA: sequncing
DEG: differentially expressing genes
FC: fold change
IHC: Immunohistochemistry

## Acknowledgements

We would like to thank Mrs. Dawn Garcia and Mrs. Korri Weldon for assistance in the performance of the single-cell RNA-seq experiments. RNA-seq experiments were conducted in the Genome Sequencing Facility (GSF) in the Greehey Children’s Cancer Research Institute (GCCRI) of UTHSCSA. The GSF facility has been constructed in part with the support from UT Health San Antonio, NIH/NCI P30 CA054174 (Cancer Center at UT Health San Antonio), NIGMS/NIH S10 Shared Instrumentation Grant Program (SIG) (S10OD021805-01 to Z.L.), and Cancer Prevention Research Institute of Texas (CPRIT) Core Facility Award (RP160732). The Flow Cytometry Shared Resource at UT Health San Antonio is supported by a grant from the National Cancer Institute to the Mays Cancer Center (P30CA054174), a grant from the Cancer Prevention and Research Institute of Texas (CPRIT) (RP210126), a grant from the National Institutes of Health (S10OD030432), and support from the Office of the Vice President for Research at UT Health San Antonio.

## Author Contributions

All authors reviewed the manuscript. A.H.H., J.M., S.A.S.: *methodology, investigation, visualization*. S.A.S., Y.Z., Z.L., A.V.T., S.R., A.N.A.: *analysis, conceptualization*. A.V.T., S.R., A.N.A.: *research design*. A.N.A.: *manuscript draft preparation.* A.V.T., S.R., A.N.A.: *resources*. A.V.T., S.R., A.N.A.: *supervision and funding acquisition*. A.V.T., S.R., A.N.A.: *final manuscript preparation*.

## Data and Material Availability

Single-cell RNA-seq data has GEO Accession number of GSE240432. It will be made available upon publication of the manuscript. Supplementary excel files show the mean gene readings for each cluster. Supplementary files are “*Non-neuronal TG clusters (1^st^ run).xlsx*” and “*Non-neuronal TG clusters (2^nd^ run).xlsx*”.

